# Reconstruction of the phase dynamics of the somitogenesis clock oscillator

**DOI:** 10.1101/743724

**Authors:** Lucas J. Morales Moya, J. Kim Dale, Philip J. Murray

## Abstract

In this study we develop a computational framework for the reconstruction of the phase dynamics of the somitogenesis clock oscillator. Our understanding of the somitogenesis clock, a developmental oscillator found in the vertebrate embryo, has been revolutionised by the development of real time reporters of clock gene expression. However, the signals obtained from the real time reporters are typically noisy, nonstationary and spatiotemporally dynamic and there are open questions with regard to how post-processing can be used to both improve the insight gained from a given experiment and to constrain theoretical models. In this study we present a methodology, which is a variant of empirical mode decomposition, that reconstructs the phase dynamics of the somitogenesis clock. After validating the methodology using synthetic datasets, we define a set of metrics that use the reconstructed phase profiles to infer biologically meaningful quantities. We perform experiments in which the signal from a real time reporter of the somitogenesis clock is recorded and reconstruct the phase dynamics. Application of the defined metrics yields results that are consistent with previous experimental observations. Moreover, we extend previous work by developing a gradient descent method for defining automated kymographs and showing that boundary conditions are non-homogeneous. By studying phase dynamics along phase gradient descent trajectories, we show that, consistent with a previous theoretical model, the oscillation frequency is inversely correlated with the phase gradient but that the coefficient is not constant in time. The proposed methodology provides a tool kit for that can be used in the analysis of future experiments and the quantitative observations can be used to guide the development of future mathematical models.

## 1 Introduction

During development of the vertebrate embryo, the head-tail axis sequentially segments into pairs of segments at regular intervals in time. Underlying this temporal periodicity is a molecular oscillator known as the segmentation clock that is characterised by waves of gene expression (Palmeirim et al., 1997; Lauschke et al., 2013; Soroldoni et al., 2014) which traverse the presomitic mesoderm (PSM) and come to rest at the boundary of future segments.

The development of new PSM culture systems (e.g. Lauschke et al., 2013; Tsiairis and Aulehla, 2016) has facilitated a much higher throughput analyses of spatiotemporal dynamics than was previously possible. Of particular relevance to this study is an *ex vivo* mPSM explant culture system in which a small, dissected part of the PSM from a live reporter mouse is cultured as a monolayer. When cultured under appropriate conditions, the mPSM explants exhibit spatiotemporal oscillations of gene expression.

The development of real-time reporters of clock gene expression in multiple vertebrate species has revolutionised the field of somitogenesis research (Soroldoni and Oates, 2011; Aulehla et al., 2008; Masamizu et al., 2006). Numerous methods have been used to process real time reporter signal so as to yield biologically meaningful inference: the time that elapses between peaks of gene expression (Lauschke et al., 2013; Webb et al., 2016; Hubaud et al., 2017); the Fourier transform (Tsiairis and Aulehla, 2016); the Wavelet Transform (Soroldoni et al., 2014; Webb et al., 2016; Hubaud et al., 2017); the Hilbert Transform (Lauschke et al., 2013); moving averages (Sonnen et al., 2018); moving average and a Savitzky-Golay filter (Matsumiya et al., 2018); representation as a harmonic oscillator (Delaune et al., 2012).

The computation of the phase of an oscillating signal allows for the relative progression through an oscillatory cycle to be quantified. In complex spatiotemporal systems, characterisation of the phase dynamics of a given signal can allow the identification of new phenomena without the need to necessarily fully understand the mechanisms (e.g. genetic components) underpinning the signal. Moreover, the inference of phase dynamics from an oscillatory signal provides a means to link experiment observations with theory.

There are numerous well-established techniques that allow the representation of temporally oscillating signals. The Fourier transform decomposes a signal into a linear sum of harmonic functions. However, for a non-stationary signal, such as a chirp signal where the frequency varies linearly time, the Fourier spectrum does not provide an intuitive representation of signal dynamics and is not particularly useful in the recovery of phase dynamics. The Wavelet Transform (Mallat, 1999), which represents a signal as a sum of predefined wavelet functions, allows for a localised description of a non-stationary signals. The results of a wavelet analysis are typically presented via a frequency spectrogram. To infer phase dynamics of the underlying signal, methods such as ridge detection are applied (e.g. Harang et al., 2012).

The application of the Hilbert transform to a signal is an alternative to template-based methods such as the Fourier and Wavelet Transforms. Here, a periodic signal is represented by

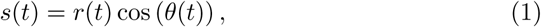

where *r*(*t*) is the instantaneous amplitude and *θ*(*t*) the instantaneous phase. Such a signal is considered to be a monochromatic signal, as it only contains a single frequency at any time point. Whilst the Hilbert transform provide provides a natural definition of oscillator phase, its application to real-world signals is complicated by the presence of trends, noise and nonstationarity (Pikovsky et al., 2001; Huang et al., 1998).

To ensure that a given signal can be studied systematically using the Hilbert Transform, Huang et al. (1998) introduced a data-driven, adaptive signal analysis technique known as Empirical Mode Decomposition (EMD). This technique, which does not require basis functions to be predefined, decomposes a signal into a set of intrinsic mode functions (IMF) that represent oscillations on distinct timescales. Each of the IMFs, which are monochromatic and can have meaningful physical/biological interpretations (Huang et al., 1998), can subsequently be analysed using the Hilbert Transform. Important limitations of EMD, which hinder its application in many practical situations, are its sensitivity to noise and mode mixing (Wu et al., 2009; Rehman et al., 2013).

The original formulation of EMD has been modified in numerous ways (Rilling et al., 2007; Rehman and Mandic, 2010, 2009; Nunes et al., 2003; Bhuiyan et al., 2008; Wu et al., 2009; Riffi et al., 2015; Wu and Huang, 2009; Rehman et al., 2013). In multivariate empirical mode decomposition (MEMD) a multi-variate signal is represented as sum of intrinsic mode functions (Rehman and Mandic, 2009). In noise assisted multivariate empirical mode decomposition (NA-MEMD) a univariate signal is processed by introducing Gaussian noise in neighbouring channels and then applying MEMD. NA-MEMD is well suited to the task of decomposing noisy time series originating from nonlinear, non-stationary oscillators.

EMD has been extended to the study of spatio-temporal signals. Bi-dimensional and tri-dimensional EMD have been developed to represent a spatiotemporal wave as a sum of spatiotemporal intrinsic mode functions with the objective being to decompose a signal into characteristic length-scales (Wu and Huang, 2009; Schmitt et al., 2014; He et al., 2017). Multidimensional Empirical Mode Decomposition (Wu et al., 2009) has been developed, in the context of image analysis, as a generalisation of EMD to two and three spatial dimensions (Nunes et al., 2003; Chen and Jeng, 2014). EMD has been combined with Principal Component Analysis or Empirical Orthogonal Functions to account for spatial structure (Park et al., 2013; Davies and James, 2014; Wu et al., 2016). Spatiotemporal MEMD is a multivariate EMD that has been used to incorporate spatial information in Brain Computer Interfaces (Park et al., 2013; Davies and James, 2014).

In this study we develop a methodology for the reconstruction of phase dynamics in an experimentally-motivated situation where a spatio-temporally oscillating signal is modulated by a time-dependent envelope. The general problem is to reconstruct the phase, *θ*(*x, t*), from a signal of the form

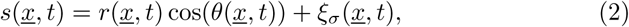

where

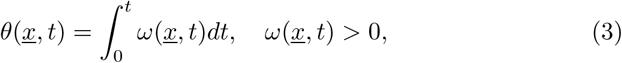

where *r*(*<u>x</u>, t*) represents the signal amplitude, *θ*(*<u>x</u>, t*) represents the phase, *ω*(*<u>x</u>, t*) represents the frequency and *ξ*_*σ*_(*<u>x</u>, t*) represents Gaussian noise of strength *σ*. The layout of the paper is as follows: in Section 2 we outline methods; in Section 3 develop and validate a methodology for phase reconstruction that is based on EMD and apply it to experimental data from a real time reporter of the somitogenesis clock; and, finally, in Section 4 we conclude with a discussion.

## 2 Methods

### 2.1 A synthetic dataset

#### 2.1.1 Model equations

We generate a synthetic dataset that exhibits the major features of real-time reporters of the somitogenesis clock in mPSM explants: an amplitude gradient, a frequency gradient, initially spatially homogeneous oscillations that are followed by the progressive loss of signal at the boundary.

Let the instantaneous phase, *θ*(*<u>x</u>, t*), be given by

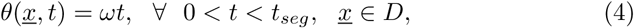

where *t*_*seg*_ represent the time at which segmentation begins. Suppose that *θ*(*<u>x</u>, t*) satisfies the partial differential equation

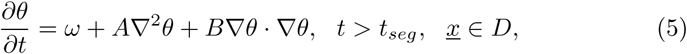

where *ω* is the natural oscillation frequency and *A* and *B* are coupling parameters (Murray et al., 2011). The domain *D* is defined to be the disk of radius *R*, i.e.

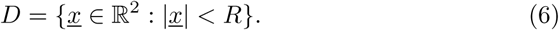

The boundary conditions are given by

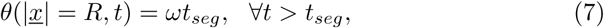

and the initial conditions are

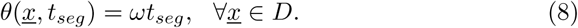

To imitate the signal decay after arrest, the amplitude is defined to be

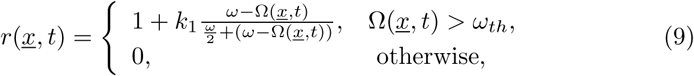

where

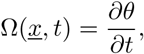

and *ω*_*th*_ represents a frequency threshold below which the amplitude is set to zero. The synthetic signal is then given by

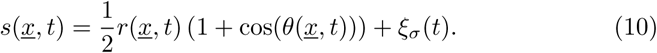

#### 2.1.2 Numerical implementation

Equations (4)–(8) were solved numerically in a Cartesian coordinate system. The spatial domain was discretised using a regular square lattice with spatial step ∆*x* = ∆*y*. Spatial operators were approximated using a central difference scheme given by

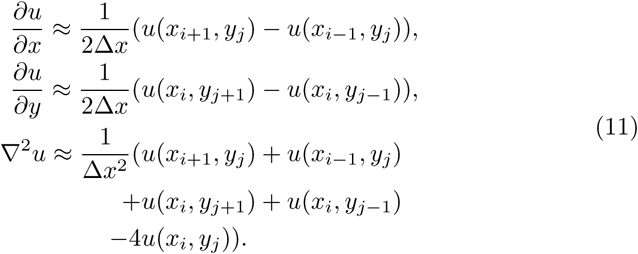

The time derivative we approximated using a Forward Euler scheme.

### 2.2 Phase reconstruction

Let *s*(x, *t*) represent a spatio-temporal oscillatory signal. The steps used to generate the reconstructed phase profile are outlined in Figure 1.

**Figure 1:**
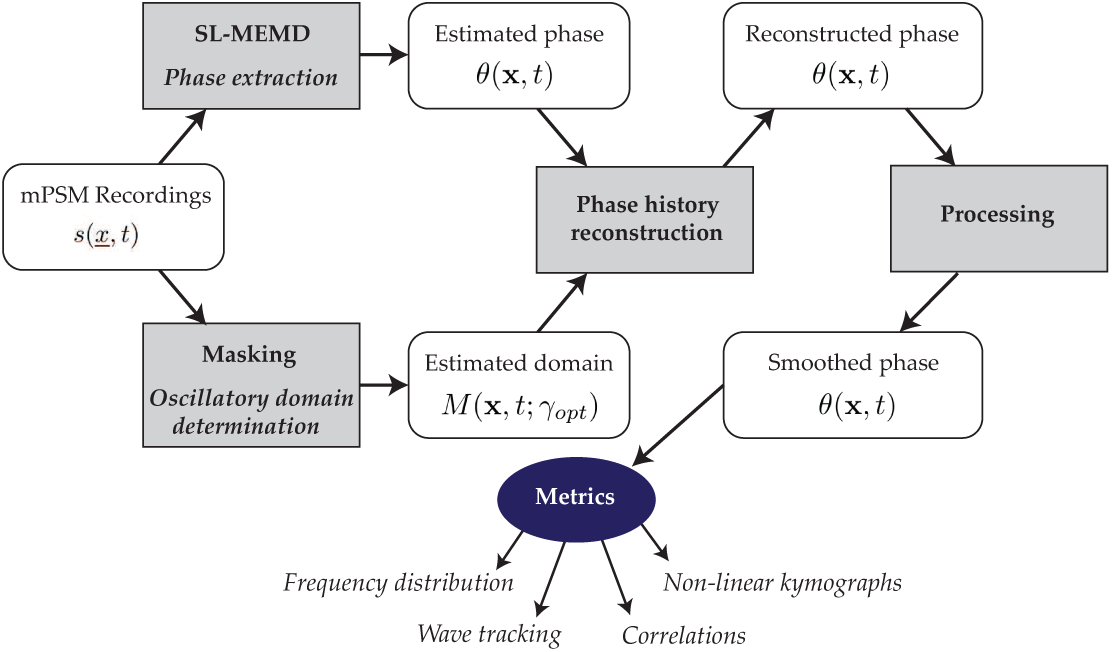
A schematic illustration of phase reconstruction.

#### 2.2.1 Multivariate Empirical Mode Decomposition (MEMD)

Consider the multivariate signal **s**(*t*) = [*x*_1_(*t*)*, x*_2_(*t*), …]. Decomposition by MEMD () proceeds as follows: Define a point set on on a (*n* − 1)-sphere (i.e. a set of direction vectors **e**_*i*_, *i* = 1,. ., *n*_*d*_). Let ŝ_*k*_(*t*) represent the *k*^*th*^ proto-IMF. Initialise by setting *k* = 1 and defining ŝ_1_(*t*) = s(*t*).

1. Project the *k*^*th*^ proto-IMF onto the *i*^*th*^ direction vector, i.e. define ŝ_*ki*_(*t*) = ŝ_*k*_(*t*) ⋅ **e**_*i*_.
2. Compute the set of local maxima of ŝ(*t*)_*ki*_, i.e. define (*t*_*kij*_, ŝ(*t*_*kij*_)), *j* = 1,. ., *n*_*ij*_.
3. Use interpolation to define the multivariate envelope, **E**_*ki*_(*t*), that pass through the local maxima of ŝ_*k*_(*t*).
4. Repeat Steps 1-3 for each direction vector.
5. Compute the mean

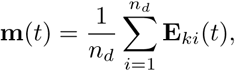

the ‘detail’

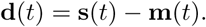

and the total variance

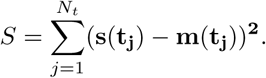
6. If *S* ≥ *S*_*stop*_, the stopping criteria are not satisfied. Define ŝ(*t*)_*k*_ = **d(t)**.
7. If *S* < *S*_*stop*_ the stopping criterion are satisfied. Define the *k*^*th*^ IMF to be **d**(*t*) and the *k* + 1^*st*^ proto-IMF to be the residual ŝ(*t*) − **d**(*t*).
8. If ŝ(*t*) has detectable extrema, repeat steps 1-7. Otherwise exit.

#### 2.2.2 Spatially localised Multivariate Empirical Mode Decomposition (SLMEMD)

Let *s*_*ij*_(*t*) represent the signal at the *ij*^*th*^ point on a regular lattice. A multivariate signal is defined with components that comprise the signal at the *ij*^*th*^ lattice point and nearest neighbours, i.e.

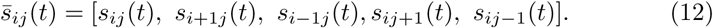

Application of Multivariate Empirical Mode Decomposition (MEMD) yields

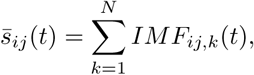

where *IMF*_*ij,k*_(*t*) represents the *k*^*th*^ intrinsic mode function at the *ij*^*th*^ lattice site and *N* is the total number of intrinsic mode functions in the decomposition.

High frequency noise is eliminated by eliminating high frequency modes, i.e.

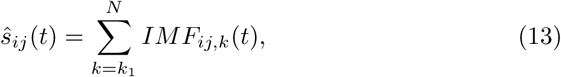

where *k*_1_ is the mode number that determines how many of the high frequency modes are filtered.

Due to (slow oscillating) trends that hinder the application of the Hilbert Transform (Pikovsky et al., 2001), the Empirical Mode Decomposition (Huang et al., 1998) is applied to the residual, i.e.

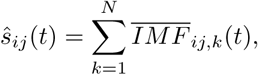

where 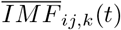 represents the *k*^*th*^ intrinsic mode function at the *ij*^*th*^ lattice site. The reconstructed signal is defined to be the first IMF, i.e.

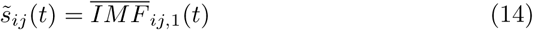

Application of the Hilbert Transform, 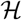, yields the analytic signal

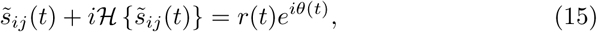

where *r*(*t*) and *θ*(*t*) are the instantaneous amplitude and phase respectively. The instantaneous frequency is computed as

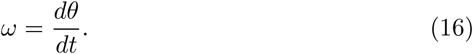

The above process is applied to all voxels in the sample.

#### 2.2.3 Identifying the oscillatory domain

To track the oscillatory region of the mPSM explant, a time-dependent mask is constructed. At a given time point, a mask is defined by thresholding the signal, i.e.

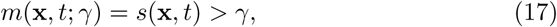

where *γ* is a threshold parameter that is optimised on a sample-by-sample basis.

To compute the boundary of the oscillatory domain, erosion is applied on the mask, i.e.

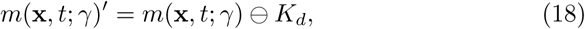

where *K*_*d*_ is a kernel of radius *R*_*d*_ and ⊖ is the erosion operator. The boundary, which is the eroded region, is given by

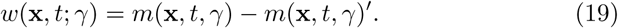

To identify the optimal value of the threshold, *W*_*γ*_ is defined to be the sum of the intensity of all voxels on the boundary, i.e.

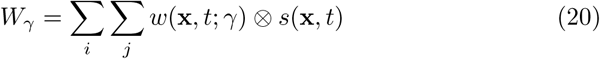

As in the experimental data oscillations are arrested after they achieve maximum amplitude, the optimal value of *γ*, *γ*_*opt*_, is defined to be

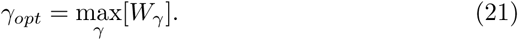

To eliminate potential holes, both dilation and erosion are applied, i.e.

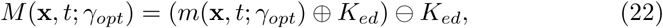

where *K*_*ed*_ is a kernel of radius *R*_*ed*_ and ⊕ is the binary dilation operator. Finally the mask is time-averaged over three time points. Applying the mask *M* (**x**, *t*; *γ*_*opt*_) to the phase yields the oscillatory region of the PSM explant, i.e.

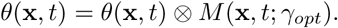

### 2.2.4 Phase on the boundary

During the segmenting phase of mPSM explants, the oscillating domain reduces in size. Additionally, the signal intensity is largest on the boundary of the tissue and there is no signal outside of the actively oscillating domain. However, the pattern has phase information (most peripheral region have experienced fewer oscillations than those in the interior).

To retain phase information, on points in space and time where the signal is being lost are identified (*<u>x</u>*^∗^, *t*^∗^), i.e.

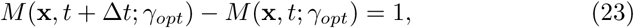

we impose the condition

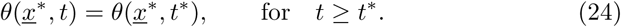

### 2.2.5 Initialisation and unwrapping of phase

The unwrapped phase is defined to be

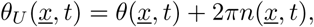

where *n*(*<u>x</u>, t*) represents the number of complete cycles that a voxel has gone through in a particular point in space and time.

In general it is not possible to infer *n*(x, 0) using the raw signal (one does not know how many cycles have elapsed at the beginning of the recording). To define an unwrapped phase in the case of mPSM explants, we use the fact that there is a maximum phase drop of 2*π* across the PSM tissue. Hence a time *t*^∗^ is identified where a region of mPSM tissue is just entering a new cycle. Hence it is approximated that

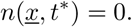

To compute the unwrapped phase profile *n*(**x, t**) for *t* > 0, ±2*π* jumps are identified using the function `unwrap` implemented in `MATLAB`).

#### 2.2.6 Smoothing of the phase profile

We compute a median filter of radius *R*_*filt*_ voxels.

#### 2.2.7 Computing sample trajectories

Let x_0_ be an initially selected point in the oscillatory domain. The phase gradient, ∇*θ*, is computed and a gradient descent algorithm is used to iteratively compute the trajectory

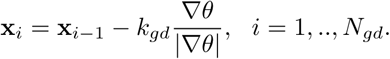

where the arc length is given by

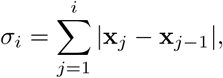

*k*_*gd*_ is the step size and *N*_*gd*_ is the maximum number of steps. The phase on the trajectory is given by

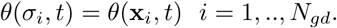

We apply a smoothing cubic spline that identifies the smoothed phase, *θ*_*S*_(*σ*_*i*_, *t*), to be the function that minimises

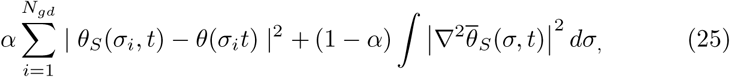

where *α* ∈ [0, 1] a regularisation parameter. The phase gradient is approximated as

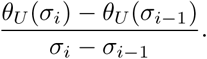

### 2.3 Metrics

#### 2.3.1 Wave tracking

To measure tissue scale periodicity, the fraction of cells that have undergone *k* cycles at a given time *t* is computed. We define the number of cycles in the actively oscillating domain to be

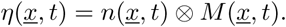

Defining the number of actively oscillating voxels at time *t* to be

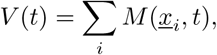

the fraction of voxels with *k* elapsed cycles is given by

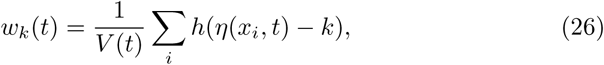

and *h*(⋅) is an indicator function given by

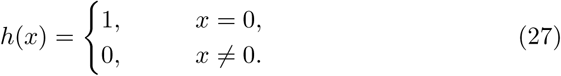

The time of the emergence of the *k*^*th*^ wave, 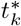, is defined to be

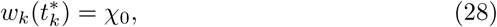

where *χ*_0_ is a constant. The tissue scale period is defined to be the time between the emergence of two consecutive waves, i.e.

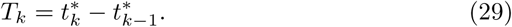

#### 2.3.2 Instantaneous frequency

Suppose that *SG*(*<u>x</u>*_*i*_,, *t*_*k*_, *ω*) is a time frequency spectrogram measured at voxel at position *<u>x</u>*_*i*_. The spectrogram of a sample is given by

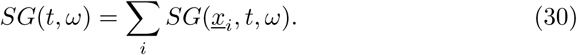

where the sum is taken over all voxels.

#### 2.3.3 Differentiation rate

The normalised differentiation rate is defined to be

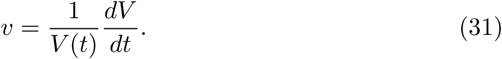

### 2.4 Experimental

#### 2.4.1 Mouse line

The *LuVeLu* mouse (*Mus musculus*) (a gift from O. Pourquie) expresses *Venus-YFP* under the control of a 2.1kb fragment of the *Lunatic Fringe* (*Lfng*) promoter. The mRNA contains the 3’UTR of *Lfng* and the protein is fused to a PEST domain, to destabilise both the RNA and the protein and ensure clear oscillations (see Aulehla et al. (2008) for further details).

#### 2.4.2 Mating procedure

Mouse E10.5 embryos were generated and the line was maintained by crossing *LuVeLu* males to stock CD1 females, as the *LuVeLu* construct is lethal in homozygotes.

#### 2.4.3 Mouse genotyping

To renew the mouse line, timed matings were performed and the litters genotyped to ensure the *LuVeLu* construct is still present. A diagnostic PCR was performed on total DNA from an ear biopsy tissue of individual animals. DNA was extracted by incubation in microLYSIS-Plus buffer (*Thistle Scientific*) through the following PCR protocol: 65°C for 15 minutes, 96°C for 2 minutes, 65°C for 4 minutes, 96°C for 1 minute, 65°C for 1 minute, 96°C for 30 seconds and 8°C until stopped.

The PCR mix was generated by adding 2*µ*l of the lysed solution to a solution containing 1.25*µ*l GoTaq™Flexy polymerase (*Promega*), 0.31 mM dNTPs (*Promega*), 1.25 mM MgCl_2_, 1 GoTaq™Flexi PCR buffer (*Promega*) and 20 pmol of each of the following four primers: (i) Ala1 (Forward), 5’-tgctgctgcccgacaaccact-3’; (ii) Ala3 (Reverse), 5-tgaagaacacgactgcccagc-3; (iii) IMR0015, 5’-caaatgttgcttgtctggtg-3; (iv) IMR0016, 5-gtcagtcgagtgcacagttt-3. Distilled water was added to the solution to reach a final volume of 20*µ*l. The following PCR protocol was used: 94°C for 2 minutes; 35 cycles of [92°C for 45 seconds, 59°C for 40 seconds, 72°C for 40 seconds]; 75°C for 5 minutes; and 4°C until stopped.

PCR samples were analysed through electrophoresis, by loading 5*µ*l of each PCR product onto a 1% agarose gel with 1:10,000 Gel Red™(*Biotium/VWR*®) and run for 20 minutes at 100 volts. The result was visualised using an UV light box. Wild type CD1 presented a single fragment of 200 bp whilst *LuVeLu*^+^*/*− presented this fragment together with a 461 bp fragment.

#### 2.4.4 *ex vivo* culture system

A 35 mm FluoroDish (*World Precission Instruments*™) with coverglass bottom was coated with a 50 *µ*g/ml fibronectin (*Sigma*) in a 100 mM sodium chloride (NaCl) solution made with double distilled water (ddH_2_O) before the dissection. The dish was incubated in the solution either for 4 hours at room temperature or overnight at 4 °C with agitation. The solution was later removed and the dish left until it was completely dried, around 30 minutes.

The dish was washed in tail bud culture medium of DMEM/F12 with no phenol red (*Gibco/Life Technologies*™) with 0.5 mM glucose (*Sigma*), 2mM L-glutamine (*Gibco/Life Technologies*™), 1% bovine serum albumin (BSA) (*Sigma*), penicillin/ streptomycin(*Gibco/Life Technologies*™) for 30 minutes at room temperature.

Embryos were harvested at E10.5 from timed matings. Individual embryos were taken from the uterine horn in sterile PBS using forceps and transferred to dissection media. To identify *luVeLu* positive embroys, tails were cut and transferred to a pre-warmed tail bud dissection media (tail bud culture media + 10 mM HEPES (*Sigma*) in a multi-well dish.

After *LuVeLu* tail identification, the tail bud was isolated from each tail posterior to the neuropore and transferred to an imaging disk, with the cut facing downwards, towards the fibronectin-coated surface.

Explants were incubated at 38.5°C, 5% CO_2_ for 1 hour to allow the explant to adhere to the fibronectin-coated surface, before live imaging. Embryos were transferred to a confocal microscope, as described in Section 2.4.5, and imaged for 24 hours at 37°C, 5% CO_2_ and ambient O_2_ levels.

#### 2.4.5 Imaging

Tail bud explants were prepared as described above. The imaging dish was transferred to a 37°C heated stage with a heated chamber at 37°C with 5% CO_2_ and ambient O_2_ of a Zeiss 710 inverted confocal microscope. Explants were imaged using a EC Plan-Neoufluar 10x/0.30 dry objective (*Zeiss*, M27). The *LuVeLu* fluorophore was excited by using a 514nm Argon laser.

Samples were scanned bi-directionally, to increase acquisition speed, and averaged 8 times per line, with a spatial resolution of 1024×1024 pixels and a temporal resolution of 15 minutes. Three optical planes were acquired at 14*µ*m intervals and starting at the plane of the cut, and moving upwards.

## 3 Results

We constructed a synthetic signal that exhibited many of the phenomena exhibited mPSM explants. The numerical solution of equations (4)–(8) exhibited a phase gradient with phase highest in the centre of the domain (see Figure 2 (a)). By computing the sine of the oscillator phase we obtain a periodic signal that is analogous to the read-out from a clock reporter (see Figure 2 (b)). To mimic the loss of signal on the peripheral boundary, we defined a dynamic spatial domain given by equation (9) (see Figure 2 (c)). Finally, we constructed an a amplitude modulated signal (equation (10)) in to mimic observations of amplitude gradient observed in mPSM explants. The goal in this section is to develop a methodology for inference of the unwrapped phase profile presented in Figure 2 (a) from the experimental-like signal presented in Figure 2 (d).

**Figure 2:**
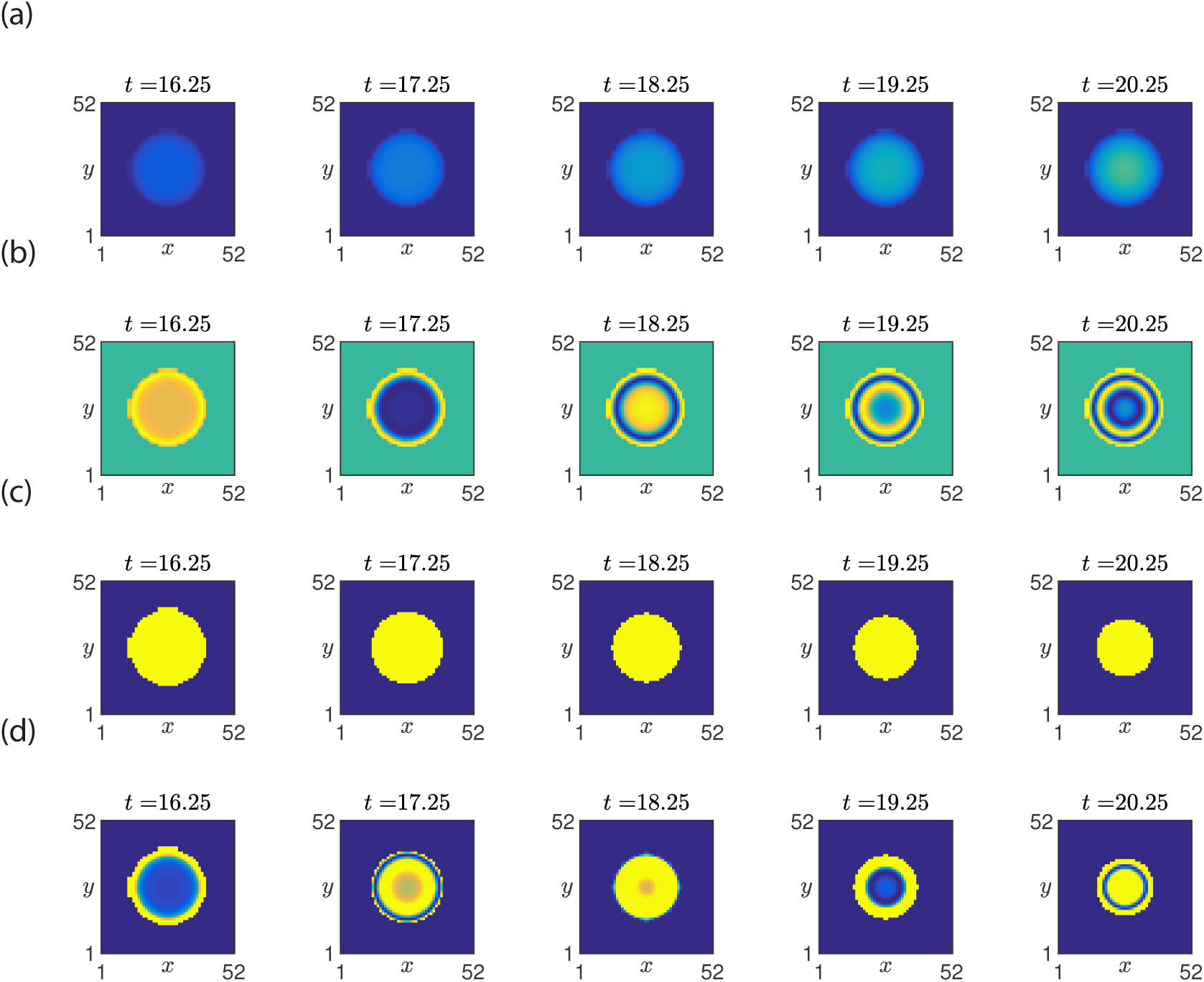
The synthetic dataset *s*(*x, y, t*). (a) Snapshots of the oscillator phase, 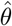. (b) Snapshots of the sine of the oscillator phase, sin(*θ*). (c) Snapshots of the actively signalling domain (equation (9)). (d) Snapshots of the synthetic signal (equation (10)). See Table 2 for parameter values.

A phase reconstruction algorithm (see Section 2.2) was developed in order to reconstruct the phase dynamics of a frequency and amplitude modulated oscillatory signal defined on a dynamic domain. The first step in the algorithm uses a novel extension of EMD that we call spatially localised empirical mode decomposition (SL-MEMD). SL-MEMD is a multivariate EMD in which a multivariate signal is defined that considers the signal in each voxel and that its nearest neighbours (see Section 2.2.2). After recovering the signal using SLMEMD (see Figure 3 (a)), a mask is defined that identifies the actively oscillating region of the signal (see Figure 3 (b)). To record the phase history in regions of the sample that have stopped oscillating, the phase is fixed in time as the boundary propagates inwards (see Figure 3 (c)). Finally, the reconstructed phase is unwrapped and smoothed using a median filter (see Figure 3 (d)).

**Table 1:** A table with parameters values used in synthetic data. c.d. cell diameter (10*µ*m).

| Parameter | Value | Definition | Unit |
| --- | --- | --- | --- |
| A | 4 | Coupling constant | c.d<sup>2</sup>h<sup>-1</sup> |
| B | 1 | Coupling constant | c.d<sup>2</sup>h<sup>-1</sup> |
| $\omega$ | 2.5 | Oscillation frequency | h<sup>-1</sup> |
| $\omega_{th}$ | 0.25 | Oscillation threshold | h<sup>-1</sup> |
| $t_{seg}$ | 16 | Segmentation time | h |
| $k_1$ | 3 | Amplitude modulation | h |
| $R$ | 20 | Domain radius | c.d. |
| $\Delta x$ | 1 | Spatial step | c.d. |
| $\Delta t$ | 0.025 | Time step | h |

**Table 2:** A table with parameters values used in phase reconstruction and metric evaluation.

| Parameter | Value | Definition |
| --- | --- | --- |
| $\chi_0$ | 0.01 | Mask threshold |
| $\alpha$ | 0.05 | Spline smoothing |
| $R_{filt}$ | 3 | Median filter radius |
| $k_{gd}$ | 1.2 | Gradient descent step size |
| $N_{gd}$ | 20 | Max steps in gradient descent |
| $\omega_{ms}$ | 0.53 | Mode selection frequency threshold |
| $R_{ed}$ | 3 | Mask erosion kernel radius |
| $R_d$ | 5-10 | Mask erosion kernel radius |
| $n_D$ | 300 | Number of direction vectors in MEMD |
| $S_{stop}$ | 0.1 | MEMD stopping criterion parameter |

**Figure 3:**
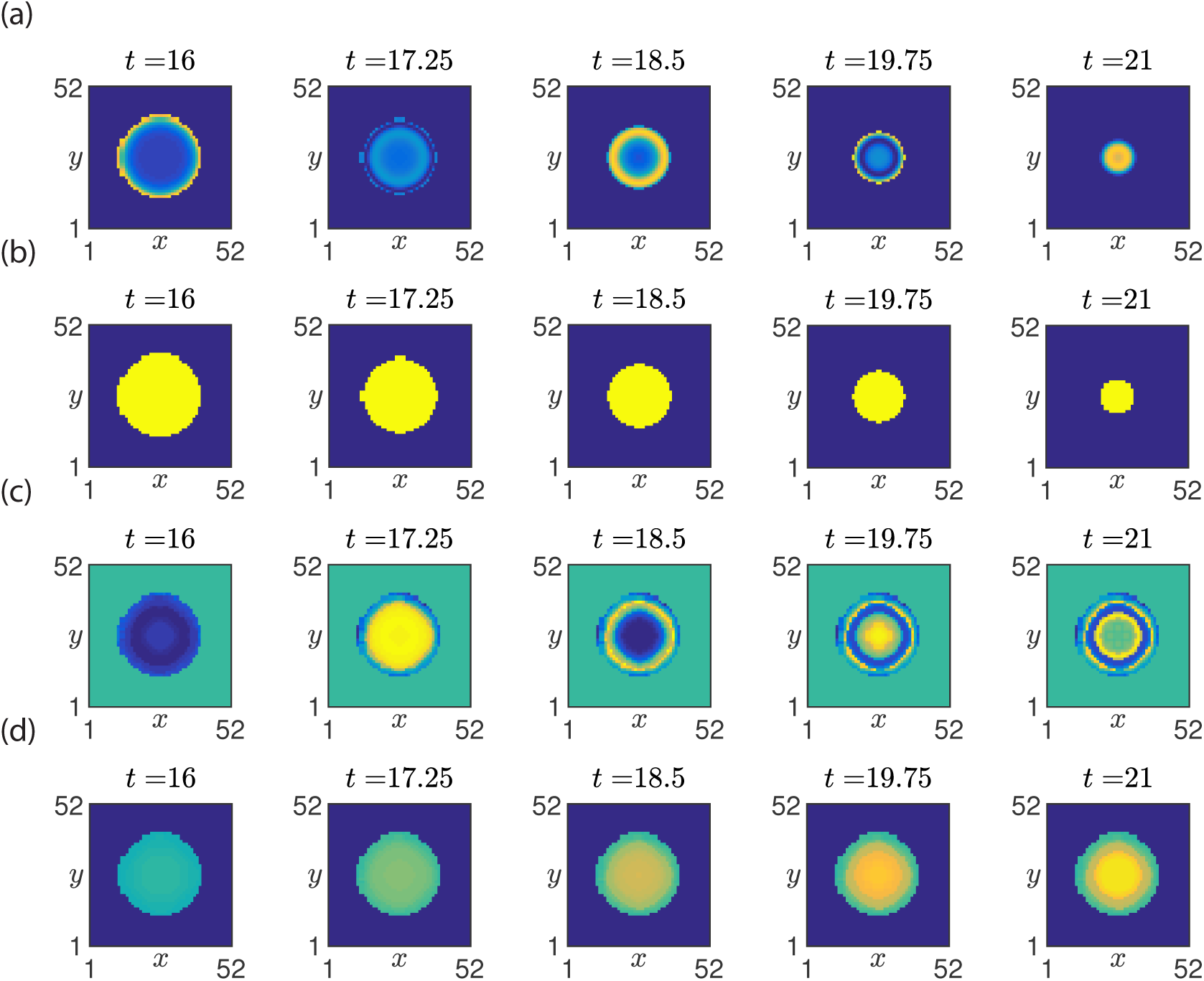
Phase reconstruction of the synthetic dataset *s*(*x, y, t*). (a) Snapshots of the synthetic signal. (b) Snapshots of the mask that defined the extent of the actively signalling domain. (c) Snapshots of the sine of the reconstructed oscillator phase, 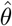. (d) Snapshots of the oscillator phase, 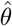. See Table 2 for parameter values.

We have developed techniques that use the reconstructed phase profile to infer biologically-meaningful quantities. To infer the tissue scale frequency, a wave tracking algorithm was developed that identifies the emergence of successive oscillatory waves (see Figure 4 (a) and Section 2.3.1). By computing the time that elapses between the emergence of successive waves (see Figure 4 (b)), the tissue scale oscillation frequency is accurately computed (see Figure 4 (c)). To describe the differentiation rate of the tissue, the normalised rate of change of area of the actively oscillating region (see Figure 4 (d)) with respect to time was computed. To describe the frequency dynamics throughout the population, we computed an instantaneous frequency spectrogram (see Figure 4 (d) and Section 2.3.2). Each of the above metrics showed excellent quantitative agreement with the ground truth computed directly from the known phase dynamics (i.e. numerical solutions of equations (4)–(8).

**Figure 4:**
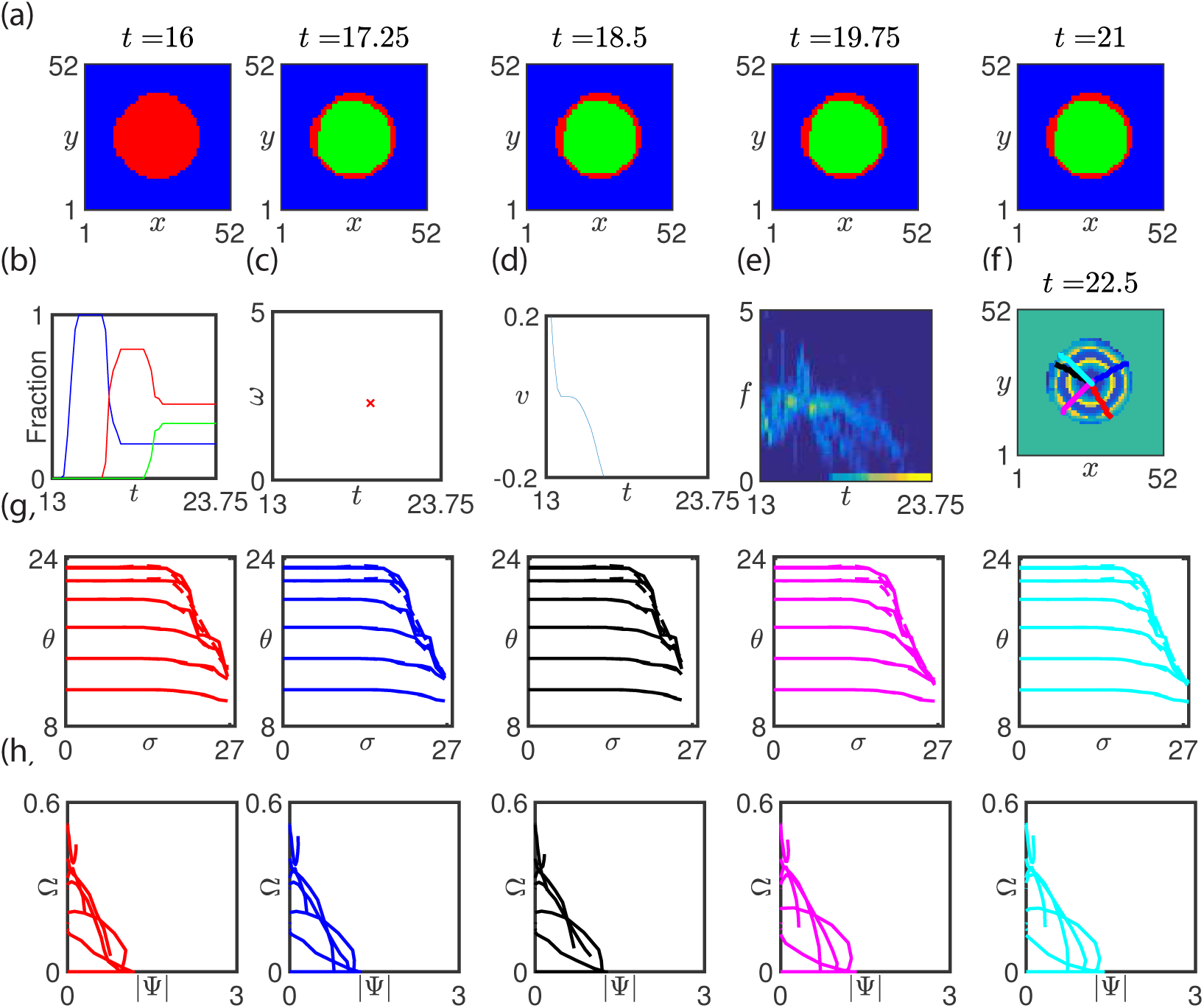
Application of metrics to reconstructed phase dynamics. (a) Snap-shots of the emergence of successive waves. (b) The area spanned by each wave is plotted against time. (c) The inter-wave frequency is plotted against time (see equation (29)). (d) The tissue differentiation rate is plotted against time (equation (31)). (e) The distribution of instantaneous frequencies is plotted against time. (f) Identification of trajectories using gradient descent. (g) Phase is plotted against arc length, *σ*, along different gradient descent trajectories. (h) The oscillation frequency is plotted against the phase gradient along different gradient descent trajectories.

To automate the generation of kymographs, we developed a gradient descent method for identifying trajectories that are locally tangential to the phase gradient (see Figure 4 (f)). In Figure 4 (e) we plot the phase dynamics as a function of arc length along five such trajectories. In Figure 4 (f) we plot the oscillation frequency against phase gradient. Note that the underlying structure of the model used to generate the synthetic data is recovered (i.e. the oscillation frequency varies inversely with the magnitude of the phase gradient). These results show that the proposed methodology is capable of describing a range of features of phase dynamics in simulated mPSM explants.

To test the proposed phase reconstruction technique a previously characterised experimental system (Lauschke et al., 2013), mPSM explants from a real time reporter of the mouse somitogenesis clock were cultured on fibronectin coated glass coverslips. We found that explants were viable and exhibited spatiotemporal oscillations of gene expression (see Figure 5 (a) and (b)). Moreover, we observed a front of high reporter activity that propagates from the periphery towards the centre of the tissue and oscillatory waves of gene expression that propagate from the centre to the periphery. However, in our hands there was significant inter-sample variability and the geometry of the sample appeared to to play a dominant role in the pattern of gene expression.

**Figure 5:**
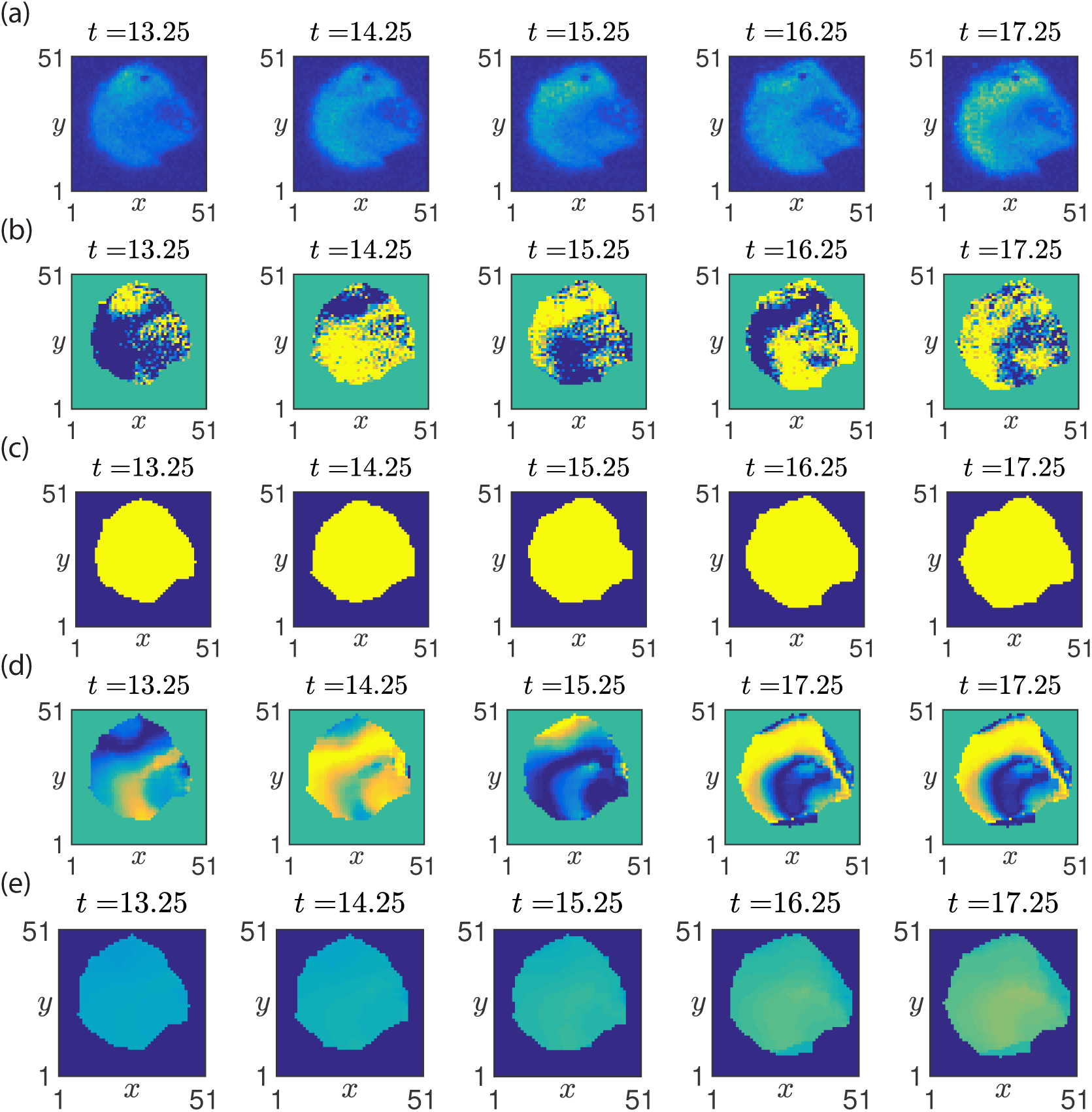
Recovered spatiotemporal dynamics from an mPSM explant. (a) Snapshots of the fluorescent signal from the LuVeLu reporter. (b) Snapshots of the fluorescent signal after application of a moving average filter. (c) Snapshots of the mask that defined the extent of the actively signalling domain. (d) Snapshots of the sine of the reconstructed oscillator phase, 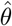. (e) Snapshots of the oscillator phase, 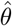. See Section 2.2 for method and Table 2 for parameter values.

To reconstruct the phase profile we applied the algorithm outline in Section 2.2. The actively oscillating region was determined (see Figure 5 (c)) and application of SLMEMD yielded the phase dynamics in the actively oscillating region of the signal. The full phase history of the sample was defined by recording the phase at the segmentation boundary (see Figure 5 (d) and (e)).

When the wave tracking algorithm (see Section 2.3.1) was applied to the reconstructed phase profile (see Figure 5 (a) – (c)), we found that the average oscillation frequency was in close agreement with that measured in a previous study (Lauschke et al., 2013). Moreover, the transition from a growing state with spatially homogeneous phase dynamics to a segmenting state with a phase gradient was identified via computation of the tissue differentiation rate (see Figure 5 (d)) and the frequency spectrogram (see Figure 5 (e)).

To explore spatial structure of the reconstructed phase profile, we used the phase gradient descent method to identify trajectories along which phase waves propagate (see Figure 6 (f)). By examining phase dynamics along these trajectories, we compute automated kymographs that allow spatio-temporal phase dynamics to be systematically explored (see Figure 6 (g)). We find that in general the boundaries of the mPSM sample are not homogeneous (at a given point in time some regions of the boundary stop oscillating whilst others do not). For trajectories that connect the centre of the explant with regions of the boundary that are still oscillating (red and blue), the phase gradient is small and the oscillation frequency approximately constant. For trajectories that connect the centre of the explant with regions of the boundary upon which oscillation had stopped, the oscillation frequency varies inversely with the magnitude of the phase gradient (see Figure 6 (h)).

**Figure 6:**
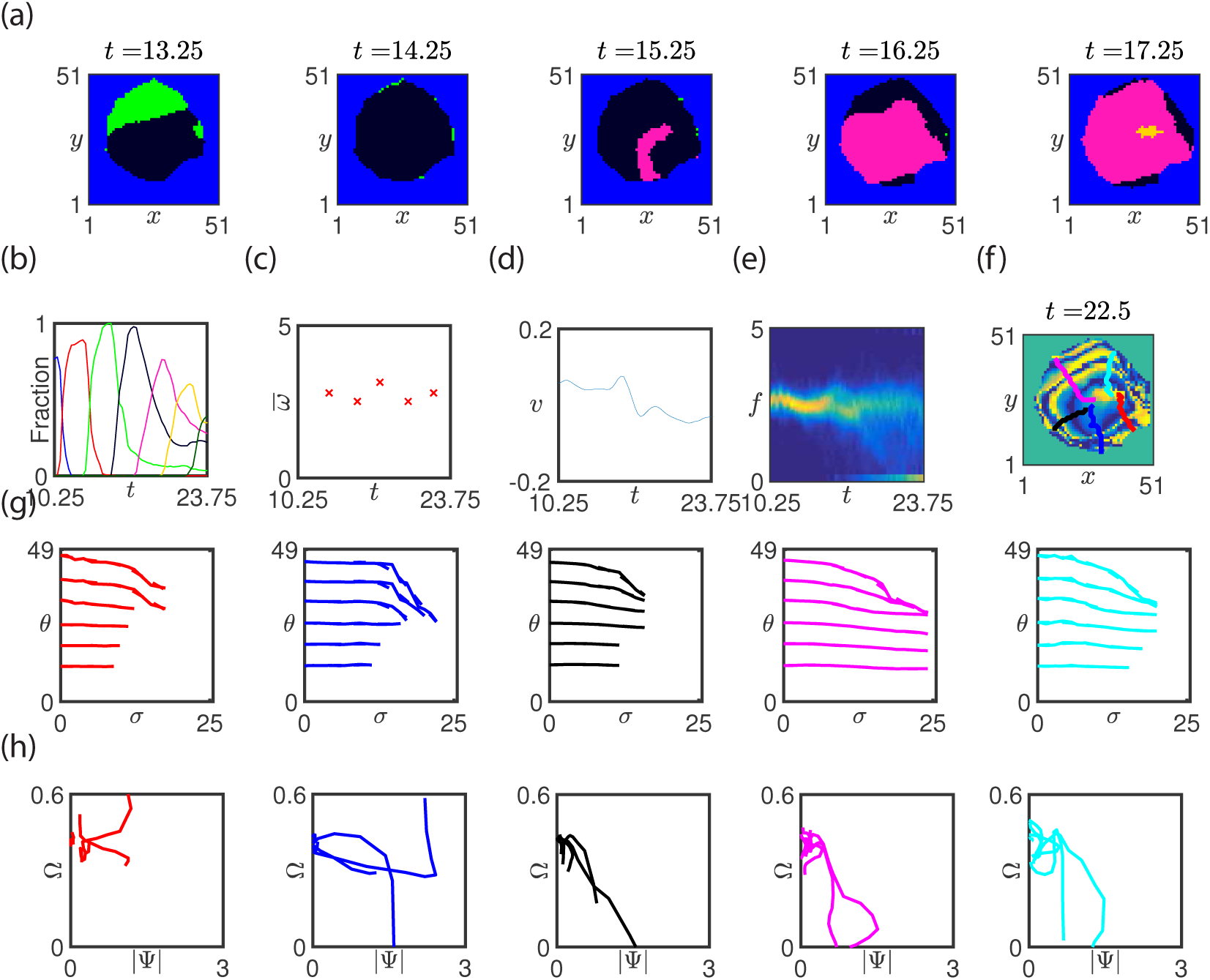
Quantitative analysis of an mPSM explant. (a) Snapshots of the emergence of successive waves. (b) The area spanned by each wave is plotted against time. (c) The inter-wave frequency is plotted against time (see equation (29)). (d) The tissue differentiation rate is plotted against time (equation (31)). (e) The distribution of instantaneous frequencies is plotted against time. (f) Identification of trajectories using gradient descent. (g) Phase is plotted against arc length, *σ*, along different gradient descent trajectories. (h) The oscillation frequency is plotted against the phase gradient along different gradient descent trajectories.

## 4 Discussion

Spatio-temporal oscillations play a crucial role in many biological systems (e.g. neural signalling, cardiac waves, calcium waves, somitogenesis). The quantification of the features of oscillations provides an information-rich means of characterising system behaviour as well as a bridge between experimental observation and theory.

The phase of an oscillator, a variable that describes relative progression through a cycle, allows the phenomenological features of its oscillatory behaviour to be described. There are numerous methods available to define phase (e.g. Fourier, Wavelet, Hilbert, EMD), each of which can yield robust phase reconstructions in certain contexts. Data-driven methods, such as EMD, are well-suited to the study of nonstationary, nonlinear signals.

During development of the vertebrate embryo, the presomitic mesoderm sequentially segments into pairs of somites at regular intervals in time. Underlying the temporal periodicity of somite formation is a molecular oscillator, known as the segmentation clock, that exhibits striking spatio-temporal patterns of gene expression. Recent advances in the development of real-time reporters of gene expression (e.g. Soroldoni and Oates, 2011; Aulehla et al., 2008; Masamizu et al., 2006) have resulted in the identification of novel phenomena, such as a Doppler effect and phase gradient scaling. There is not yet a consensus on which methods are optimal for quantifying phenomenology of the spatio-temporal dynamics.

In this study we have developed a phase reconstruction methodology based on empirical model decomposition that allows inference of phase dynamics on a dynamic spatial domain. The methodology uses a variant of EMD, which we call SLMEMD, to infer phase dynamics. Moreover, we develop a set of metrics that when applied to the reconstructed phase profile yield the quantification of biologically meaningful variables.

In order to validate the phase reconstruction methodology, we defined a synthetic dataset that mimicked many known features of segmentation clock dynamics. We showed that the ground truth phase could be accurately reconstructed using the proposed methodology and that a range of biologically relevant quantities (e.g. tissue scale period, instantaneous frequency distribution, geometrical features) can be accurately inferred from the reconstructed phase profile.

To determine if the phase construction methodology could be applied to experimental measurements from a real time reporter of the segmentation clock, we performed a series of experiments in which mPSM explants from the LuVeLu reporter mouse were cultured as monolayers on fibronectin-coated glass slides. We inferred the phase dynamics in the mPSM sample using the phase reconstruction methodology and, consistent with previous observations, found: (i) two distinct phases of behaviour (expanding and segmenting); (ii) outwardly propagating oscillatory waves of gene expression; and (iii) an inwardly propagating wavefront that defines a peripheral limit of the oscillatory domain. Application of the developed metrics yielded quantitative results that are in good agreement with the previous observations.

We have proposed SLMEMD, a novel variant on EMD that is applicable to locally synchronised spatio-temporally oscillating systems. The defining property of SL-MEMD, which is that the signal in neighbouring voxels is used in the inference of phase at a given voxel, increases the robustness of phase inference compared with EMD but does not require the tuning of noise strength as is the case with NAMEMD. We note that we have reproduced our results using NAMEMD and have not found significant deviation between the methods for the signals that we have considered.

EMD and its derivative methods are empirical. To deal with this property, we have developed synthetic datasets that approximate the experimental data from which we would like to infer phase dynamics. Whilst we have explored perturbations round these datasets, we note that aspects of the methodology may need to be optimised for sufficiently different problems.

One of the advantages of the phase reconstruction approach adopted in this study is that the phase dynamics are reconstructed in the full spatial domain. Compared with kymograph analyses, which are frequently used in the literature and require the arbitrary specification of a spatial axis upon which to analyse the signal, the proposed approach allows one to quantify the evolving geometry of phase profile.

A limitation of the current study is that individual cells are not tracked. In principle, artefacts in the phase dynamics could be induced by cell motion. In the mPSM explant analysed in this study, we considered only the segmenting phase (*t* > 16*h*) during which cell motion is greatly reduced.

An important caveat with this work is that the experimentally measured signal is periodic. Hence at given instant in time one sees a readout of gene expression but not the absolute number of cycles that have elapsed. This issue is problematic in the definition of the unwrapped phase as it is impossible to infer the absolute phase the reporter signal. However, mPSM explants exhibit approximately spatially homogeneous patterns of gene expression at early times, a property that is used to define the initial condition for the unwrapped phase.

In this study we have developed a methodology for reconstructing phase dynamics in mPSM explants. In future work we will use the phase reconstruction techniques to investigate the response of mPSM explants to chemical perturbation and we will explore the extent to which low dimensional mathematical models can be used to reproduce observed phenomenology.

